# Turnover shapes evolution of birth and death rates

**DOI:** 10.1101/2022.07.11.499527

**Authors:** Teemu Kuosmanen, Simo Särkkä, Ville Mustonen

## Abstract

Population turnover, a key trait shaped by the organism’s life history strategy, plays an important role in eco-evolutionary dynamics by fixing the timescale for individual birth and death events as well as in determining the level of demographic stochasticity related to growth. Yet, the standard theory of population genetics, and the models heavily used in the related data analysis, have largely ignored the role of turnover. Here we propose a reformulation of population genetics starting from the first principles of birth and death and show that the role of turnover is evolutionarily important. We derive a general stochastic differential equation for the frequency dynamics of competing birth-death processes and determine the appropriate turnover corrections for the essential results regarding fixation, establishment, and substitution of mutants. Our results reveal how both the absolute and relative turnover rates influence evolution. We further describe a deterministic turnover selection, the turnover flux, which operates in small populations. Finally, we analyse the evolution of mean turnover and show how it explains the key eco-evolutionary mechanisms underlying demographic transitions. In conclusion, our results explicitly show how competing life-history strategies, demographic stochasticity, ecological feedback, and evolution are inseparably intertwined, thus calling for a unified theory development starting from the underlying mechanisms of birth and death.

## INTRODUCTION

Population genetics focuses on the interplay of different evolutionary forces and its rich theory lies at the heart of quantitative description of evolution [1]. Its models provide the theoretical backbone for drawing insight from the vast data sets of the post-genomic era and have been successfully applied to explain diverse evolutionary phenomena ranging from understanding adaptation and speciation to elucidating cancer evolution, drug resistance and human demographic history [2–6]. Whereas big data biology has resulted in recent breakthroughs in quantifying evolutionary forces, e.g., the identification of distinct mutational processes active across cancers [7], the central theoretical foundation of evolution starting from first principles has remained largely unchanged and elusive.

The ubiquity of genomic data has led to a highly gene-centric view of evolution, where information gained from the large-scale analysis of genomes are seen as the main causes and putative explanations for evolutionary change. Population genetics itself enjoys a special status within theoretical biology in its view of defining selection at the level of competing alleles and emphasis on random processes which can provide sufficient explanation for evolution even in the absence of natural selection. This has resulted in debates regarding the role and importance of selection to molecular, genetic, and even phenotypic evolution [8, 9], as well as in various other specific contexts, including cancer evolution [10].

Irrespective of whether signals of selection are found from the genomes or not, no amount of high-throughput data can change the fact that natural selection ultimately results from the almost tautological consequence of differential birth and death rates of the *phenotypes* present in the population. Likewise genetic drift results from the intrinsic demographic stochasticity necessarily present in finite populations, driven by the random fluctuations between the underlying birth and death events. Remarkably, what often tends to be omitted both analysis-wise, but sometimes even conceptually, is that the primary causes of evolution have therefore fundamentally an *ecological*, and not genetic, basis [11].

Recently, even mutational processes have been linked to be at least partly modulated by the environment, as more evidence of adaptive mutability has been accumulated in unicellular and cancer cell populations [12, 13]. Epigenetic inheritance, phenotypic plasticity and other strictly non-genetic mechanisms which can create phenotypic diversity subject to selection further confound eco-evolutionary dynamics [14], thus calling for integration rather than separation of ecology and evolution. Therefore, it follows that unlike the central dogma of molecular biology would suggest, evolution does not obey the one-directional information flow originating from genes to higher levels of biological hierarchy, but instead acts in the opposite direction, where the selection at the higher-order biological entities in fact constrains the potential evolution that can occur independently at the lower levels.

Even though powerful in explaining the evolution of allele frequencies and other genomic data, the standard theory of population genetics largely fails to account for the tangled genotype-phenotype map [15], or even acknowledge the distinct spaces in which evolution operates, thus disconnecting itself from the underlying mechanisms of birth and death. Consequently, classical models of population genetics seldom capture the population dynamics seen in real populations, thereby obscuring the complicated relationship between natural selection and evolution. On the other hand, while other established fields of theoretical biology have predominantly studied the rich mechanisms of deterministic selection within ecological communities, they rarely specify how the phenotypes of interest arise or are inherited nor is it often clear how the effects of stochastic evolution could be incorporated, or even understood qualitatively (see e.g. [16]). Further ambiguities that result from the fact that different modelling frameworks often apply different proxies for “fitness” [17, 18] seem to create additional obstacles to unification of evolutionary theory.

In search for a unified understanding of evolution grounded on a solid mechanistic foundation, birth-death processes omnipresent across all scales of biology have been proposed as a viable alternative [18]. We agree that this is a promising direction and explicitly show here how birth-death processes open the path forward both conceptually and also in terms of mechanistic modelling of evolution. Instead of building ecology on top of some widely used evolutionary models, such as the Wright-Fisher model [19] or the Moran model [20, 21], here we propose a reformulation of population genetics starting from the underlying birth and deaths events and their respective rates, which can describe arbitrary population dynamics. This way, as we will show, the impressive machinery of population genetics can be harnessed and extended to ecologically more realistic assumptions and systems while still allowing analytical insights to the stochastic evolution of novel mutants.

There exists a large body of literature covering the subject of fixation of beneficial mutations under various assumptions [22–25], which also includes life-history [26–29] and birth-death-like models [30–32]. However, most of the previous work has been done under the limiting assumptions that mutations change either only the birth rate (fecundity-mutants) or the death rate (generation-time mutants) while the other trait remains constant [24]. Alternatively, Parsons *et al*. consider the so-called quasi-neutrality case where the ratio of the birth and death rates is fixed [31]. By allowing the birth and death rates remain completely general, we show how these restrictive assumptions have concealed the surprisingly prominent role the turnover rate - here defined as the sum of birth and death rates - plays in evolution.

By deriving a stochastic differential equation for the frequency dynamics of competing birth-death processes, we show how the classical Wright-Fisher diffusion [33] can be recovered as a special limit where all the turnover rates scale to one. After arguing why this limit is unrealistic in natural populations, we proceed to rederive the quintessential results concerning the fixation, establishment, and substitution dynamics of mutants, in effect providing the appropriate turnover corrections to them. Our analytical results show how these key quantities substantially depend on the turnover rates. We further describe a deterministic turnover selection term, the turnover flux, which selects for lower turnover mutants analogous to Darwinian selection, in small populations. We then turn to ask how the turnover itself, as a quantitative trait, evolves and derive an equation governing the evolution of the mean turnover rate in a population.

## RESULTS

### From competing birth-death processes to population genetics

Consider a population of replicators with distinct subtypes *i* = 1, …, *d*, each with some intrinsic birth and death rates 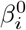 and 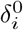, respectively. Let us assume that these intrinsic rates are both heritable and mutable, and thus subject to evolution via genetic innovation, and that they determine the growth trait 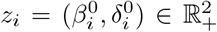 by which the replicator subtypes would initially grow exponentially in a pristine environment, in the absence of any ecological interactions between the replicators. (In the case of more complex multicellular organisms, the growth trait would rather form an entire (possibly age- or size-dependent) life-history strategy 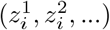 but here we are primarily interested in constant life-history strategies, widely applicable in modelling cell and microbial populations.) A fundamental question in biology is then to explain and make predictions about the stochastic evolution of these growth phenotypes while factoring in the potentially complex ecological interactions which regulate the underlying, intrinsic rates, and thus lead to the actually observed, effective birth and death rates by which the real population dynamics unfolds. This distinction between the intrinsic and the effective rates is important because only the intrinsic rates are evolvable by mutation while it is the effective rates by which the evolutionary forces of natural selection and genetic drift are actually experienced (see Fig. 1).

**FIG. 1.**
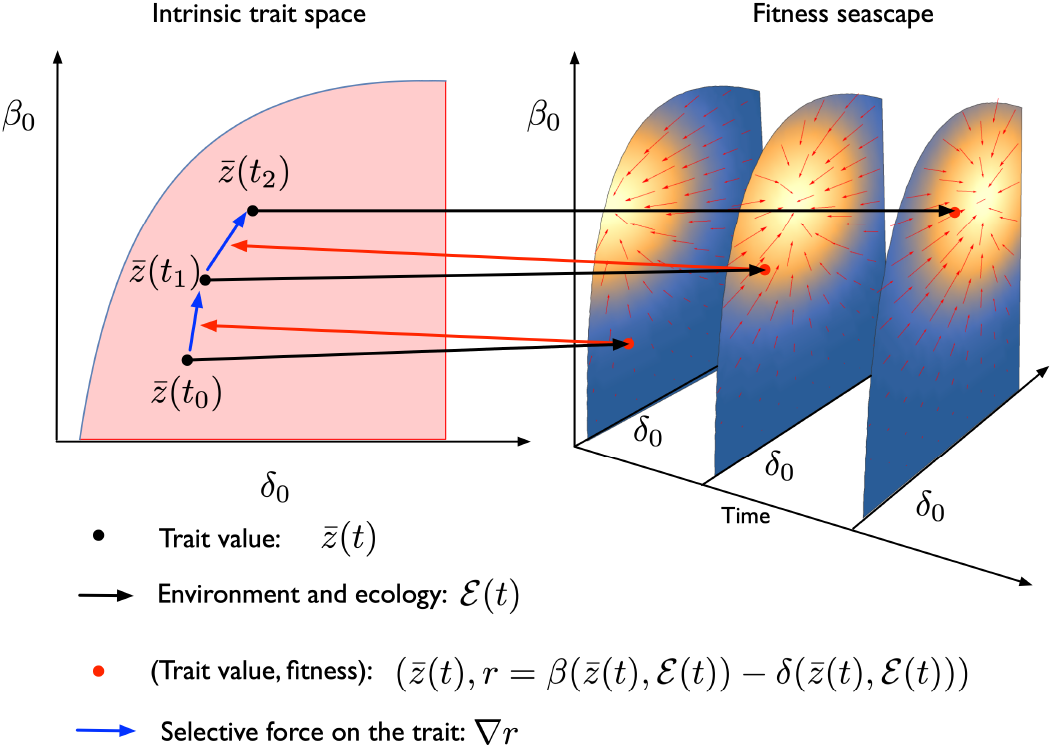
Eco-evolutionary feedback loop. Mutation generates diversity to the intrinsic birth and death rates present in the population, and these competing life-history strategies in turn translate to environmental feedback *ℰ*, creating a mapping (black arrows) to the space of effective birth and death rates. The difference of the effective birth and death rates determines the instantaneous fitness landscape (visualized by the heatmap on the effective planes) which maps back to the intrinsic space (red arrows) via selection of intrinsic traits, causing the mean trait 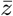 to evolve along the gradient of the fitness landscape 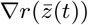 (blue arrows). The new population state once again feeds back to the environmental variable *ℰ*, thereby dynamically changing the fitness landscape via density- and frequency-dependent interactions. Such time-dependent fitness landscapes are called fitness seascapes [34]. The focus of the present study is to investigate how the described evolutionary dynamics changes both qualitatively and quantitatively in the presence of demographic stochasticity, where also the intrinsic volatility related to the expected growth rate given by the fitness landscape is taken into account. As we will show, also the turnover rate *λ*, described by the sum of the effective birth and death rates, plays a fundamental role in evolution, shaping the probability distribution of the evolutionary trajectories.

In what follows, we understand the effective birth and death rates to be arbitrary functions of the trait vector *z* = (*z*_1_, …, *z*_*i*_, …, *z*_*d*_), total population size *N* = Σ_*i*_ *X*_*i*_ (where *X*_*i*_ denotes the population size of subtype *i*), frequency vector 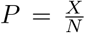, and time *t* to accommodate all possible density- and frequency-dependent interactions as well as environmental stochasticity. To simplify notation, we introduce environmental feedback variables (*ℰ*_*β*_(*t*), *ℰ*_*δ*_(*t*)) = *f*(*t, z*(*t*), *N* (*t*), *P* (*t*)) that the system’s ecology determines via the function *f*. Now, the effective rates for subtype *i* can be written as functions *β*(*z*_*i*_, *ℰ*_*β*_) and *δ*(*z*_*i*_, *ℰ*_*δ*_), where the corresponding environmental feedback variable regulates the respective intrinsic rate either additively or multiplicatively. As we will later show, this property of the ecological feedback, too, will prove to be important.

We begin by introducing a general stochastic differential equation (SDE) which describes how the frequencies of the different subtypes change in the population (for full derivation starting from the discrete-time birth-death process, see Methods-section). First, let us denote the difference between the effective birth and death rates as *r*_*i*_ := *β*(*z*_*i*_, *ℰ*_*β*_) − *δ*(*z*_*i*_, *ℰ*_*δ*_), which gives the expected growth rate of subtype *i* at the given environment (and is thus time-dependent, and not constant), essentially corresponding to the fitness-generating function discussed in [35]. Furthermore, let us denote the sum of the effective birth and death rates as *λ*_*i*_ := *β*(*z*_*i*_, *ℰ*_*β*_) + *δ*(*z*_*i*_, *ℰ*_*δ*_), which we define as the *turnover rate* of subtype *i* (also time-dependent).

Now, the population size *X*_*i*_ of subtype *i* is a stochastic birth-death process approximated by (22) (see Methods), and we can apply Ito’s lemma (see e.g. [36]) to recover the corresponding SDE for the frequency dynamics of the strain of interest. This yields

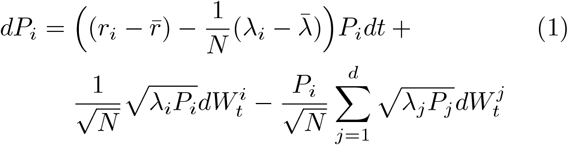

where 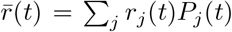 is the current mean growth rate in the population, 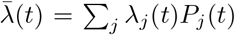 is the current mean turnover rate, 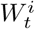 are independent Wiener processes, and the total population size *N* is stochastic with the associated equation

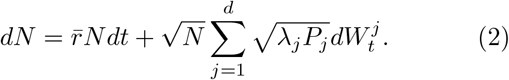

When looking at the deterministic part of (1), we immediately recognize its first term 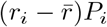 as the well-known *replicator equation* (see e.g. [37]), which simply states that the strains, which grow faster (slower) than the population as whole, increase (decrease) in frequency. We notice that the frequency dynamics of arbitrary population dynamics in principle reduces to the replicator dynamics in the limit *N* → ∞, thus providing a highly general and simple description for the effects of natural selection in large populations. However, in the reality of finite populations, we see that the role of turnover stands out and intriguingly another analogous turnover term 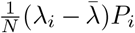 appears even to the deterministic part, acting on top of the classical Darwinian selection for faster growth. We call this term the *turnover flux* because it results solely from the probabilistic flow caused by the demographic stochasticity related to the birth and death events, in turn favouring strains with lower turnover and smaller stochastic fluctuations.

We next analyse the important special case of *d* = 2 to get the much-studied mutant-resident model. We can express the mutant frequency *p* = *P*_1_ using a single Wiener process term as

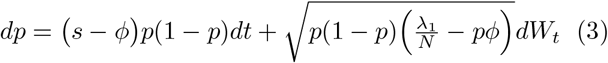

where we have now defined the classical selection coefficient as *s* = *r*_1_−*r*_2_ and the turnover flux 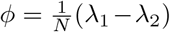. This comes at the cost of modifying the noise terms in the associated *N*-dynamics (see Methods and (29)). If interested only in the statistical properties of the mutant frequency, similar weakly equivalent SDE can also be found for the general equation (see (30)).

We note that the Wright-Fisher diffusion [33] can be recovered from (3) in the limit (*λ*_1_, *λ*_2_) → (1, 1), and it follows that the Wright-Fisher model thus formally corresponds to a specific birth-death process, where all the competing variants share the same precise turnover scaled to one. This widespread assumption built in the standard theory of population genetics is not necessary and is unlikely to be valid in natural populations. Fortunately, it can be easily remedied by switching to analyse the more general equations presented above. Indeed, we next take the mutant-resident model as our starting point and rederive the essential classical results regarding fixation, establishment and substitution of mutants, showcasing the effects of differential turnover on these quantities.

### Turnover modulates mutant fixation probabilities causing turnover bias

Given equation (3), we can write down the associated Fokker-Planck equation, which in principle allows us to solve the time-evolution of the probability distribution for the mutant frequency. From this we can calculate the fixation probability *u*(*p*), which gives the ultimate probability for the event that the mutant starting from initial frequency *p* invades and fully replaces the current resident.

Following population genetics tradition, we begin by studying the simplest case of the mutant-resident model with constant growth and turnover rates. By neglecting the ecological feedback, this is equivalent to studying mutant fixation in the pure birth-death model and allows us to quantify the effects of differential turnover rates before moving on to analyse how the ecological feedback changes the evolutionary dynamics. To do this we simply modify the Kimura equation [22] to calculate the fixation probabilities analytically. Indeed, we find that the fixation probability for a mutant originating from a single individual is given by

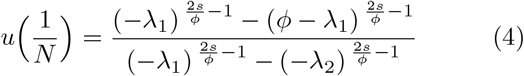

where all the rates are still the same as their intrinsic counterparts as explained. Although the expression (4) is somewhat complicated, the important limit cases turn out to be very simple and intuitive. First, for a neutral mutant we find that

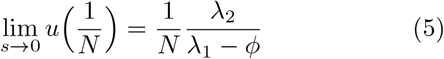

and for the case of large populations

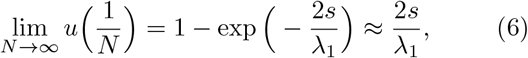

where the last well-known approximation works well when *s* is small. These turnover corrections to the classical results elegantly reveal many important insights that the Wright-Fisher limit has concealed. In small populations the classical expectation for a neutral mutant fails due to the turnover flux and also the relative turnover differences become important. In large populations, where the turnover flux formally vanishes, the mutant’s own absolute turnover rate remains evolutionarily significant because all mutations are necessarily initially rare and thus very much subject to stochastic extinctions. Consequently, it follows generally that the fixation probability for given mutant with arbitrary classical selection coefficient is always a monotonically decreasing function of its turnover.

We can next study subsequent mutant fixation events by analysing substitution dynamics. In the mutant-resident model, we can express the probability of finding the population in a monomorphic state by using a two-state approximation (which is accurate in a mutation limited case) with the appropriate substitution rates (see e.g. [38]):

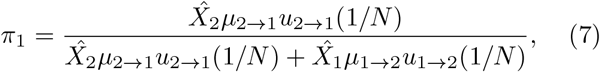

where *µ*_*i*_:s denote the mutation rates between the strains, *u*_*i*_:s the corresponding fixation probabilities, and 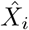:s the respective equilibrium population sizes. Now, assuming unbiased mutations and similar resident equilibria, the monomorphic state probabilities exhibit a clear *turnover bias* whereby the population, on average, is more likely to be found at the lower than higher turnover state.

### Birth-death framework allows studying fixation and substitution of mutants in ecological models

The classical theory has already been extended in special cases by various authors to include more complicated scenarios, where some of the ecologically unrealistic assumptions have been relaxed (see e.g. [23–25]). In general, we believe that studying the rich mechanisms of natural selection within ecological communities probabilistically, beyond the standard deterministic analysis based on the invasion fitness, holds the key in unifying different areas of theoretical biology. The birth-death framework allows naturally to extend the presented analysis to models with ecological feedback simply by substituting in the effective birth and death rates. However, depending on their functional forms, fully general analytical results may not always be achievable.

For the sake of concreteness and as a proof-of-principle, we now move on to fix a specific ecological model and use it for the remainder of our study to showcase the effects of turnover and ecological feedback on the evolutionary dynamics. By incorporating multiplicative density-dependent feedback on the births, additive density- and frequency-dependent feedback on the deaths, and a scalar parameter *θ* ∈ [0, 1] which determines how the total feedback is divided between the births (*θ* = 0 only births) and deaths (*θ* = 1 only deaths), the patch model defined by the effective birth and death rate equations (31) provides a sufficiently rich yet analytically tractable case study for benchmarking theoretical results to stochastic simulations of the underlying birth-death processes rooted in generalizable principles of ecological feedback (see Methods for more details).

Figure 2 shows how the analytically obtained, approximative fixation probabilities (38) closely match to the stochastic simulations of the patch model: Fig. 2A shows that the fixation probability of mutant with arbitrary selection coefficient is a monotonically decreasing function of its turnover, as already seen in the case of a pure birth-death process in Eq. (6). Fig. 2B demonstrates how the turnover flux operates and how substantially turnover modulates the fixation probabilities *u*(*p*) as a function of the mutant frequency *p*. Finally, Fig. 2C verifies the expectation of turnover bias seen in the monomorphic state probabilities.

**FIG. 2.**
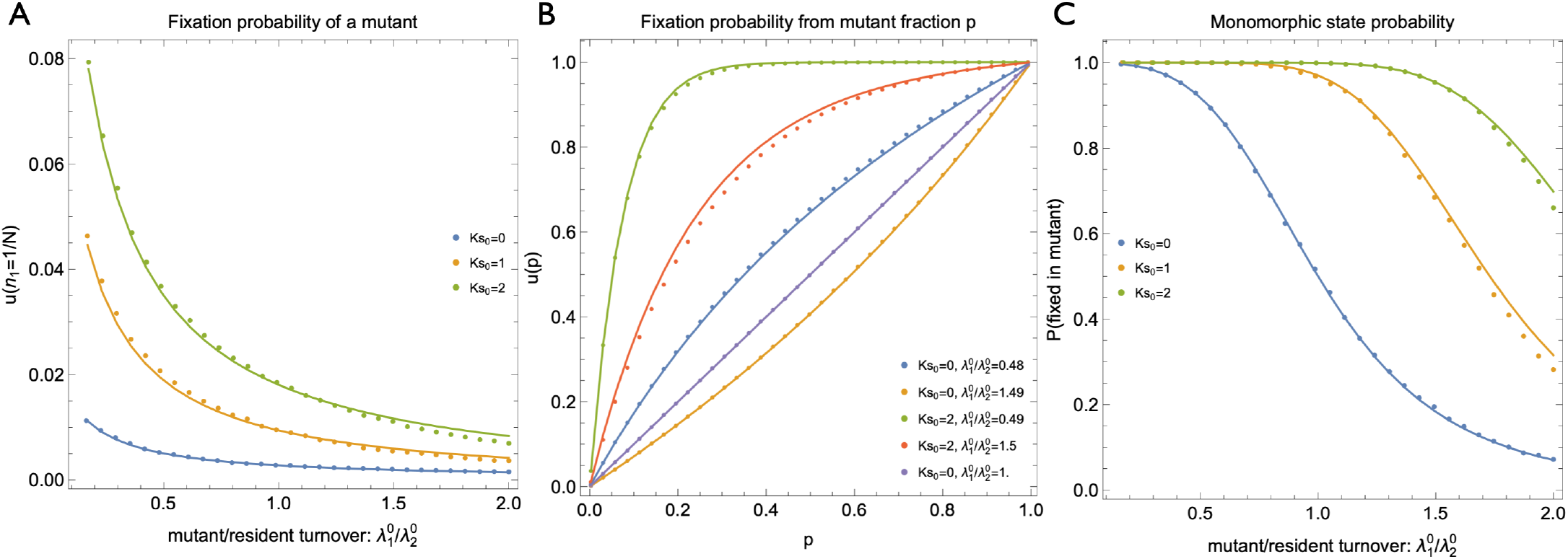
Turnover modulates fixation and substitution dynamics. **A** A single mutant was introduced to a resident population and the ecological dynamics was simulated using the patch model (with patch size *K*) until the mutant either went extinct or became fixed. The analytically solved fixation probabilities (solid lines) closely match to the simulation results (dots) obtained from the stochastic birth-death simulations, and reveal that for any given scaled selection coefficient *Ks*_0_ the fixation probability decreases monotonically as the mutant’s turnover increases. **B** Classical theory predicts that the fixation probability of a neutral mutant equals its current frequency in the population (purple solid line). However, in small populations this prediction fails due to the turnover flux, which induces selection even for classically neutral mutants, favouring the types with lower turnover (blue) over higher turnover (yellow). Similarly, lower (green) and higher (red) turnover can either promote or hamper classical growth rate selection respectively. **C** Assuming an equal mutation rate between the low and high turnover subtypes, the probability of finding the population in monomorphic state exhibits a turnover bias, whereby the population spends more time on average in the low turnover state.

We note that although similar observations related to demographic stochasticity have been made also in the literature in other contexts (see [27] and [31] for Fig. 2A and 2B respectively), we believe that our general treatment, derived from the underlying principles of birth and death, together with the turnover corrections to the classical fixation results provides further clarity on the subject and indeed shows the importance of turnover in evolution, a result that has not been fully embraced despite the intuitive notions presented in earlier works.

### Establishment threshold quantifies the escape from stochastic extinction and is modulated by turnover

In more complex ecological settings, mutant invasion will rarely lead to substitution and a monomorphic state, but rather successful invasion implies that the mutant will be able to coexist as part of the diverse ecological community. Yet, even in the case that the invasion fitness is positive, the mutant must still first escape from the stochastic extinction before the deterministic dynamics will take over and dominate genetic drift. This process of mutant establishment is a key driving mechanism by which the turnover rate affects the ultimate fixation probabilities, and conversely understanding the process establishment is conceptually attractive because it allows us to extend the key qualitative and quantitative predictions about novel mutants also in situations where a complete selective sweep is unlikely to occur.

The transition from the initial stochastic regime to the deterministic phase is known in the literature as the *establishment threshold* [39, 40], essentially representing an evolutionary equivalent of an Allee threshold in population dynamics. Rouzine *et al*. [39] derive the establishment threshold as the mutant frequency

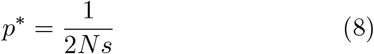

above which selection is assumed to dominate drift. However, the precise mathematical characterization and interpretation of the establishment threshold is difficult firstly, because there does not exist any objective threshold above which the mutant could not go extinct (as seen also in Fig. 2B), and secondly because the stochastic extinction risk is a property determined both by the mutant’s growth trait (birth and death rates) as well as the properties of the community it is introduced to. Therefore, given our previous analysis, it seems likely that differential turnover rates will play a key role also in defining the establishment threshold, an element missing from the previous analyses.

Next, we present a simple way to characterize mutant establishment in the introduced general birth-death framework, inspired by the so-called time-average growth rate [41]. Geometric Brownian motion exhibits an interesting phenomenon, whereby the ensemble average grows exponentially with the expected growth rate while any single trajectory will go ultimately extinct [42], as is technically the case with any population. What this essentially means is that a large enough collection of new mutants will collectively grow according to their invasion fitness irrespective of what the level stochasticity is, but the fate of any single mutant can differ considerably, and is strongly influenced by the volatility it experiences. Because many biological systems of interest are mutation-limited, looking at what happens to the typical mutant is therefore more informative than the ensemble average of such mutations.

We can analyse how the median trajectory of the mutant frequency evolves by taking the log-transformation of the frequency dynamics by applying Ito’s lemma to Eq. (3) (see [41]). This gives us the following equation for the mutant’s log-frequency (see Methods)

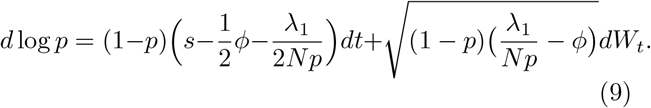

Now, we immediately see that the expected log-frequency change can initially be negative even when the mutant has a high positive invasion fitness, reflecting the fact that the deterministic selection kicks in only when the mutant has managed to grow sufficiently large. Because the establishment threshold quantifies precisely how far the given mutant must drift by luck before it can trust to enjoy positive deterministic selection, we can thus define the establishment threshold as the mutant frequency at which the typical mutant starts to grow, or equivalently when the expected log-frequency change becomes positive. Equating the deterministic part of (9) to zero thus gives

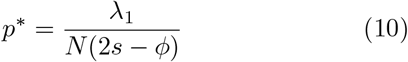

from which we again notice how the classical establishment threshold (8) is recovered in the Wright-Fisher limit.

We note that when the ecological feedback modifies the effective rates, the establishment threshold technically dynamically changes too. However, as the establishment typically occurs at relatively low mutant frequencies, when the mutant has only negligible effect on the environment, we can readily treat the establishment threshold as constant, evaluated at the initial resident feedback. Indeed, we can see from Fig. 3 how the analytically calculated establishment threshold captures the qualitatively different patterns of establishment below and above the threshold, further showing how mutants with lower turnover can establish more easily.

**FIG. 3.**
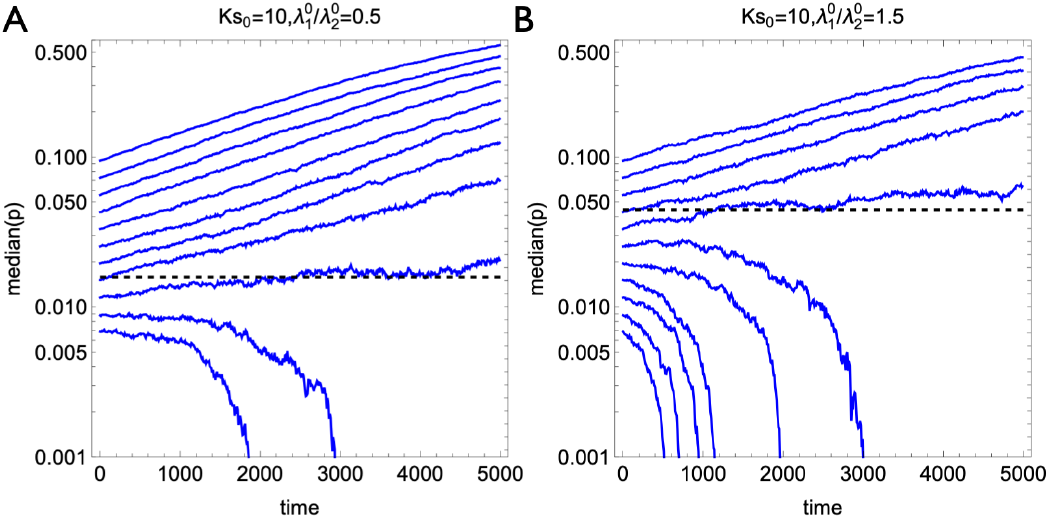
Mutant establishment threshold is modified by turnover. Low (A-panel) and high (B-panel) turnover mutants with identical positive invasion fitness were introduced to a resident population at varying initial frequencies both above and below the analytically calculated establishment threshold (dashed lines). Above the establishment threshold the frequency of a typical mutant will increase exponentially while below the establishment threshold the typical mutant will be lost from the population. Low turnover mutants establish at lower population frequencies than high turnover ones.

### Life-history evolution is determined by the variance in birth and death rates

So far we have studied the impact of turnover to the fates of mutants and showed how the demographic stochasticity determined by the turnover rates influences the process of mutant establishment and fixation, thereby causing turnover bias where mutations associated with lower turnover are more likely to be selected in the long run. When looking at the grand picture of evolution across the tree of life, we indeed witness a highly repeatable pattern of increase in organismal complexity and corresponding slowdown of life-history strategies. Next we attempt to close the circle and continue to ask how the turnover rate itself, as an quantitative trait, evolves.

When looking at the frequency dynamics, much attention has been devoted to understanding the effects that result from the stochasticity of the total population size *N*. However, we note that, critically, also the mean fitness 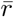 and turnover 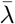 are both stochastic even though they can be easily computed from the definitions at every time step once the (*p, N*)-dynamics has been fixed. However, could we say something about their time-evolution generally without fixing some particular ecological model?

Indeed, we can now simply apply Ito’s lemma again to the definitions and so recover well-defined SDEs for the mean fitness and turnover given the general equation (1) for the frequency dynamics. If we first look at the mean fitness, we notice that we can write its deterministic part, still under completely general treatment, in surprisingly compact form as

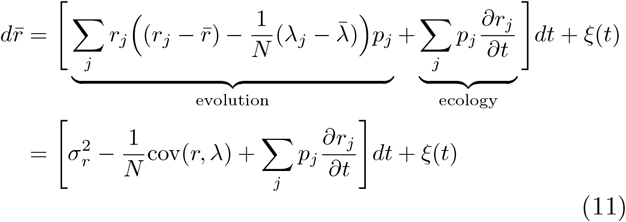

where we have now suppressed the more complicated noise term to a stochastic process *ξ*(*t*). When looking at the deterministic part of this equation, we recognize that the change in mean fitness can be decomposed, essentially in the spirit of the well-known Price-equation (see e.g. [43]), to the effects of evolution changing the frequencies of the different rates in the population, and the effects of ecology, which in contrast may dynamically change the effective rates themselves in response. Intriguingly, the evolutionary part of this equation, irrespective of the acting ecological feedback, further simplifies to being just the variance of effective growth rates 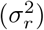 present in the population plus additional covariance term from the turnover flux.

If we neglect the turnover flux term, we recover exactly the fundamental theorem formulated in [44], which shows that the contribution of natural selection on the change of mean fitness is mediated solely by the **variance in fitness** (and not, for example, directly by genetic or phenotypic diversity as previously thought), and that in the absence of ecological feedback, selection always has a non-negative contribution to mean fitness (because variance cannot be negative). However, even though natural selection will always increase the frequency of the “fitter” individuals, this positive contribution to the mean fitness may be more than offset by the potential detrimental effect from the ecological feedback that results from the new environmental state to which the population evolves under selection. Thus, even if natural selection always increases mean fitness in relative terms, the mean fitness may still in some cases decrease in absolute terms, and in extreme cases potentially lead to evolutionary suicide [45].

Importantly, we can now repeat the exercise for the mean turnover to see that

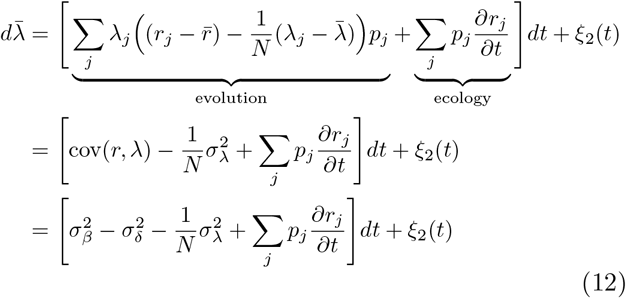

where in the last line we have further simplified the covariance term by noting that the growth and turnover rates can be further decomposed to the effective birth and death rates, which conveniently eliminates the covariances altogether. If we again first only focus on the effects of natural selection and ignore the turnover flux term, we can see that it is the variance in the effective birth and death rates which determines, at the most fundamental level, whether organisms evolve to faster or slower life-histories. Because selective advantage can only result either via mutations with higher birth or lower death, selection acting on variance in birth rates necessarily increases turnover, whereas selection acting on variance in death rates decreases turnover. Therefore, it is the difference between these two variances that dictates the direction of life-history evolution, as shown by the selection part of (12).

We can now use this equation to explain the key eco-evolutionary mechanisms underlying demographic changes, especially demographic transitions, whereby a permanent decrease in the effective turnover rate is achieved. Understanding the mechanisms leading to demographic transitions is of interest firstly, because its close connection to the well-documented changes seen in human populations [46], but also because of the highly repeatable evolutionary pattern of “complexifying force” observed across the tree of life, whereby different life-forms seem to predictably evolve ever higher organismal complexity, which in turn is associated with lower turnover.

By introducing birth and death rate mutants to the patch model in different environments by varying the parameter *θ*, we can track how evolutionary innovations (whether genetic or cultural) spread and lead to positive net growth until the ecological feedback eventually re-equilibrates the population size to its new carrying capacity. Figure 4 shows the four extreme cases where pure birth or death mutants were introduced to the extreme environments with either solely birth- or death-mediated regulation, and displays the resulting evolutionary and demographic patterns. Whereas the evolutionary dynamics looks almost identical irrespective of the type of the mutant or the ecological feedback, the resulting demographic change in the turnover is different in each case. The change in the mean turnover rate can be decomposed to the evolutionary and ecological parts as shown by Eq. (12), and decrease in turnover (a demographic transition) can only be observed in the case where an innovation, which decreases the intrinsic death rate, spreads to an environment where the ecological feedback acts by decreasing the births. Even if the intrinsic traits under selection are associated with lower (intrinsic) turnover, a demographic transition nevertheless fails to occur if the consequential ecological feedback reverses the preceding evolutionary change in turnover by equilibrating the population size with externally mediated sources of mortality (e.g. predation or parasitism). This directly connects our results to the observations in literature [47, 48], where the role of extrinsic mortality in life-history evolution has on one hand been emphasized and deemed important, but on the other remained somewhat unclear. Indeed, we can now dive even deeper in disentangling the complementary effects of selection, genetics and ecological feedback on the direction of life-history evolution by investigating how the (genetically-codified) intrinsic turnover rate evolves in the population.

**FIG. 4.**
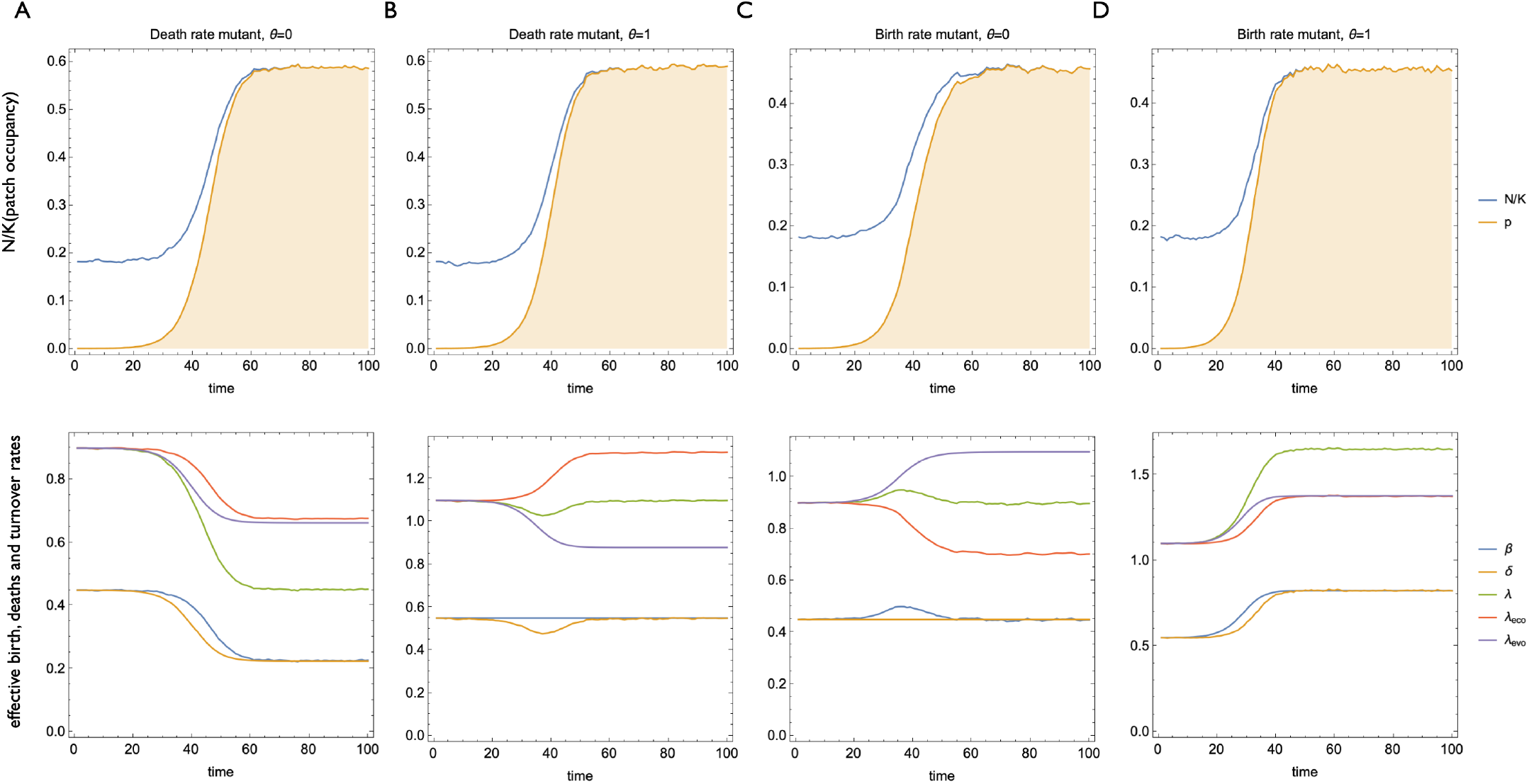
The interplay of eco-evolutionary dynamics underlies demographic transitions. All selectable mutations can be divided to innovations which either decrease the intrinsic death rate (“death rate mutants”, panels **A**,**B**) or increase the intrinsic birth rate (“birth rate mutants”, panels **C**,**D**). Upon successful mutant invasion (upper panels) also the mean (effective) birth and death rates will change accordingly (lower panels), and thus in both cases lead to positive net growth until the ecological feedback equilibrates the population size to its new carrying capacity. The ecological dynamics will then modulate the effective birth and death rates independently of the preceding evolutionary dynamics, and can either sustain (panels **A**,**D**) or reverse (panels **B**,**C**) the evolutionary change in the effective birth and death rates, leading to four distinct patterns of demographic change depicted in the lower panels. The changes in the mean turnover rate (green lines) can be decomposed to the effects of selection (purple lines) and ecology (red lines) as shown by equation (12).

### Only multiplicatively acting ecological feedback has evolutionary significance

The power of the derived fundamental equation (12) lies in the fact that we can now use the properties of variance to broadly categorize how different modes of ecological feedback induce selection for different life-history strategies also at the level of intrinsic traits. Because the variance is invariant with respect to the location parameter, it follows that only **multiplicative feedback** can have evolutionary significance. Indeed, we can now write one last equation for the *trait evolution* of the intrinsic turnover rate *λ*^0^ by considering all the four possible different combinations of ecological feedback, where both the birth and death regulation can act either additively or multiplicatively. Because the intrinsic rates are fixed, the ecological part of (12) must be zero, and we get

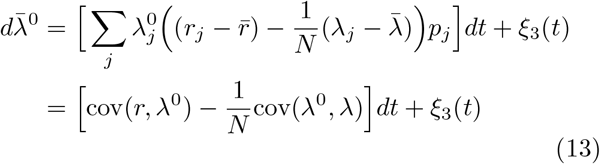

where the driving covariance term can now be simplified as

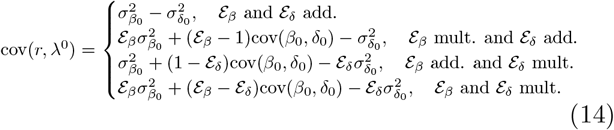

where 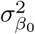 and 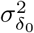 now represent the variances in the intrinsic birth and death rates, respectively. This equation now allows us to explicitly see how different modes of ecological feedback (additive or multiplicative) factor into the selection of either slower or faster turnover.

As a concrete example, let us again look at the patch model and simulate evolutionary trajectories while assuming an unbiased Gaussian mutation kernel for the intrinsic birth and death rates as well as a concave trade-off curve for the feasible trait space (see Methods). From the patch model equations (31) we can see that the birth rate regulation is multiplicative and the death rate regulation is additive, as is common in extrinsic mortality generally, and we thus get the following selection gradient for the mean turnover dynamics in the patch model (neglecting the turnover flux term)

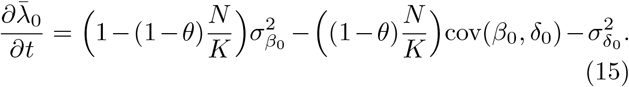

We can now use this equation to predict how the evolutionary trajectories will look like as a function of the parameter *θ*. We first note that when no life-history trade-offs are present in the interior of the trait space, the variance terms must be equal 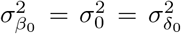 and the covariance term must vanish (when the mutational kernel is assumed to be non-biased). This simplifies the mean turnover equation in the interior trait space to the form

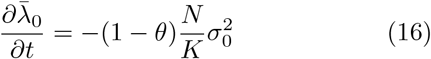

readily predicting that the mean intrinsic turnover rate will decrease whenever *θ* < 1, that is, when there is any feedback that decreases the births. This is exactly what we also observe in the simulations displayed in Fig. 5 when looking at the mean evolutionary trajectories across the *λ*^0^-contours. However, once the evolutionary trajectory hits the trade-off curve, or more generally any other (developmental) constraints in the trait space, the variance terms will no longer be equal and thus alter the turnover dynamics together with the additional covariance term. Therefore, (14) reveals how the effects of selection show up very simply as the difference in the variances of the intrinsic rates, whereas the effects of genetics and its background factor into the variance and covariance terms, which can then be further adjusted by the multiplicatively, but not additively, acting ecological feedback.

**FIG. 5.**
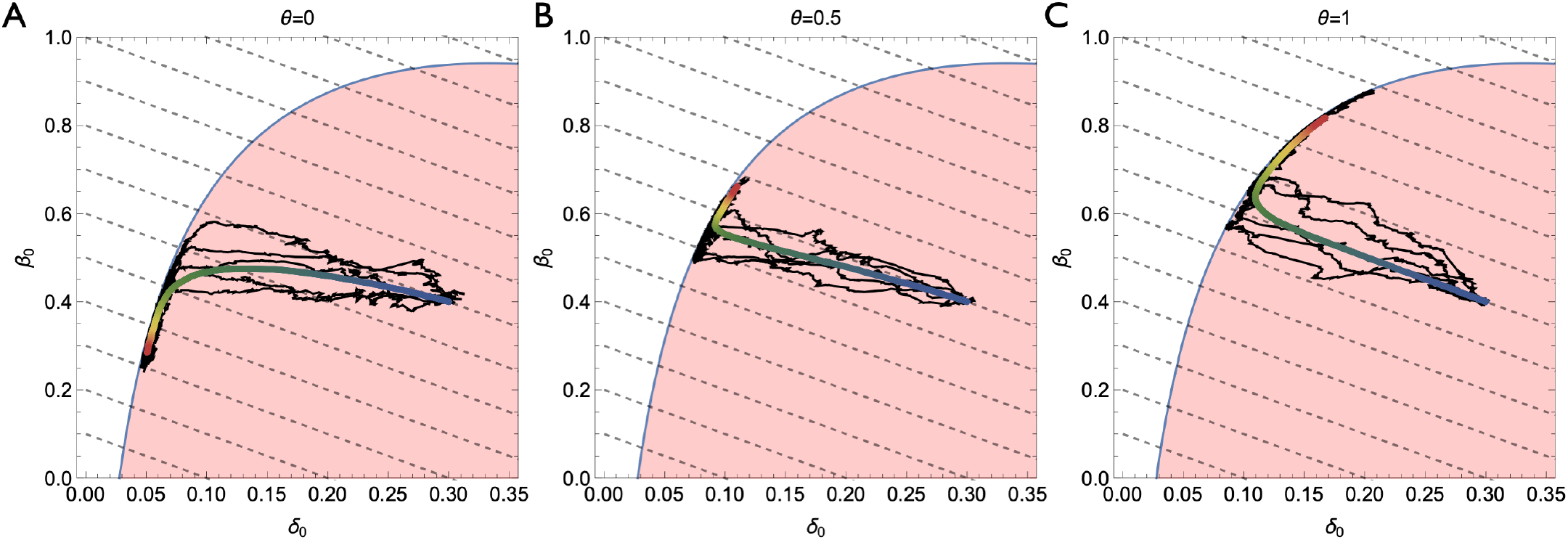
Ecological feedback mechanisms determine long-term evolutionary trajectories. Evolution of intrinsic birth and death rates in the patch model under different modes of population regulation (panels **A-C**) when the feasible trait space is assumed to be restricted by a concave trade-off curve (blue solid line). The thick lines represent the mean traits over time (shown by the color scale starting from blue and ending to red), together with five individual trajectories (black). Turnover iso-contours are shown in black dashed lines. **A** Regulation by birth-limiting environments (*θ* = 0) select for lower turnover. **C** Environments which increase effective death rate (*θ* = 1) lead to evolution of faster life-histories and higher turnover. **B** A combined regulation by births and deaths (*θ* = 0.5).

## DISCUSSION

Birth and death are ubiquitous events across all levels of biological systems, and all aspects of evolution can ultimately be traced back on how the different evolutionary forces shape organisms’ reproduction and survival. Importantly, differential birth and death rates not only determine the expected growth rate (and therefore selection) via their difference, but also set the intrinsic volatility related to this growth via their sum, the turnover rate. Even though the field of population genetics is well-known for taking the effects of stochasticity seriously, the fact that new mutants may differ not only in terms of their expected growth rate, but also in terms of their turnover, has been largely ignored. While numerous studies have investigated and identified demographic stochasticity [25, 29] and life-history strategies [27] as important factors and explored their evolutionary consequences, the concept of turnover has remained fuzzy and the standard theory of population genetics and the models used in practice say very little about the role of turnover in evolution generally. Given the surprisingly simple yet fundamental equation (12) that governs the evolution of the mean turnover, defined as the sum of birth and death rates, we indeed believe that this concept is central to evolution, as evidenced also by the fixation and establishment results exemplified in Figs. 2 and 3.

Instead of building on top of the widely used classical evolutionary models, such as the Wright-Fisher model or the Moran model, here we attempted to derive the correct evolutionary model directly from the first principles of birth and death while factoring in the multitude of ecological mechanisms that may regulate the effective birth and death rates. By doing so, we showed how the classical results based on the Wright-Fisher diffusion can be recovered as a special case where all the turnover rates scale to unity, and further demonstrated how taking this limit has in fact concealed the prominent role turnover plays in evolution. We then verified this mathematical expectation and benchmarked the analytical results with stochastic simulations of the birth-death processes using the patch model as a case-study, which is rich enough to sufficiently display the key ecological feedback mechanisms yet simple enough to preserve a degree of generality and allow meaningful comparison to the classical results.

After understanding how turnover affects the fixation probabilities of mutants, we moved on to dissect the process of mutant establishment. By refining the concept of establishment threshold [39], we showed how the initial escape from stochastic extinction can be quantified in a model-free way, showing how it is both the absolute and relative turnover rates which impact mutants’ evolutionary success. Therefore, the effects of differential turnover cannot be sensibly eliminated using scaling tricks related to “effective population sizes” because this would be unique for every mutant, and not population, thus rendering the concept practically useless. However, using the derived establishment threshold, the key qualitative role of turnover can be extended also to more complex ecological communities, where a complete fixation cannot be expected, but instead the question of coexistence arises. Given the simple and highly interpretable formula (10), one particularly interesting area for future research would be linking the here discussed establishment threshold to the concept of minimal infectious dose and understanding its determinants [49].

Finally, we investigated the long-term consequences of turnover to life-history evolution. By calculating the monomorphic state probabilities for subsequent substitution dynamics, we observed a turnover bias, where at any given time the population is on average more likely to be found at the lower turnover state. When combined with any potential macroevolutionary forces, the turnover bias alone may help to explain the apparent slowdown of life-histories across the tree of life in long evolutionary times even in the absence of direct selection for slower turnover. Extending the presented analysis of the most fundamental eco-evolutionary mechanisms underlying demographic transitions (see Fig. 4) to include e.g., age-dependent birth and death rates, may provide an interesting future direction to properly investigate such speculations and could also be of potential interest when modelling human demography [50] and cultural evolution.

The presented results are also of particular relevance to rapidly evolving cell populations, such as cancer and microbial cell communities, which can exhibit diverse life-history strategies and great turnover differences, ranging from hyperproliferative cells to complete quiescence. Indeed, the role of turnover seems to be evident especially in the context of cancer evolution [51, 52], where detailed understanding of the mechanisms of birth and death can further elucidate patterns of tumor progression and waiting times to cancer as well as guide optimal treatment decisions [53, 54]. However, in order to fully leverage the utility of the presented theory and test its predictions, it is necessary to be able to experimentally access the birth and death rates, and not only their difference. This key message of our work calls for more experimental work much in the spirit of [55], where we currently have surprisingly little knowledge about the varying rates of turnover and death even in different healthy tissues. This point is no less relevant even for a purely modelling perspective, where more emphasis ought to be put on the clear distinction between mechanisms of birth and death, and how evolution and ecology respectively affect them. This is particularly important for individual-based simulations where it is important to realize that such implicit decisions regarding how e.g., the mutations and density-regulation are implemented, can in fact cause unwanted and unrecognized evolutionary pressure towards certain phenotypes.

## METHODS

### Deriving the frequency dynamics

The population dynamics of subtype *i* with effective birth and death rates *β*_*i*_ := *β*(*z*_*i*_, *ℰ*_*β*_) and *δ*_*i*_ := *δ*(*z*_*i*_, *ℰ*_*δ*_) can be written using a discrete-time equation

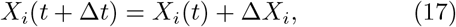

where

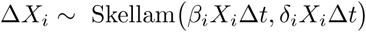

with the mean and variance of

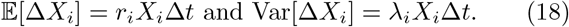

As the process is a sum of Skellam-distributed random variables (which in turn are differences of Poisson processes), we can form a coarse-grained approximation of the model by replacing the increment with a Gaussian increment with the same first two moments

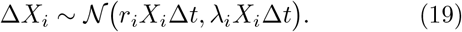

We can now introduce a Gaussian random variable

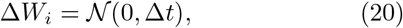

which then lets us to write (17) as

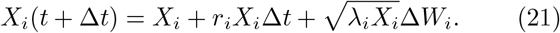

In the limit Δ*t* → 0 this becomes the stochastic differential equation

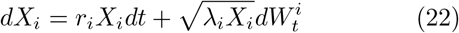

for which the discretized model can be seen as Euler– Maruyama approximation (additionally see [56] where an equation of this form is also derived).

We can now apply Ito’s lemma to function 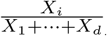 in order to get the corresponding SDE for the frequency dynamics of subtype *i*, which straightforwardly gives equations (1)-(2). Importantly, note that the equations hold for arbitrary time-dependent growth and turnover rates, because when applying Ito’s lemma the derivatives are only calculated with respect to frequency of the strain of interest, thus allowing the study of completely general population dynamics.

In the case *d* = 2, we can define *p* = *P*_1_, in which case we have 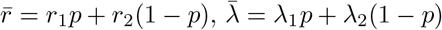. Thus, we get the following special cases of (1)-(2):

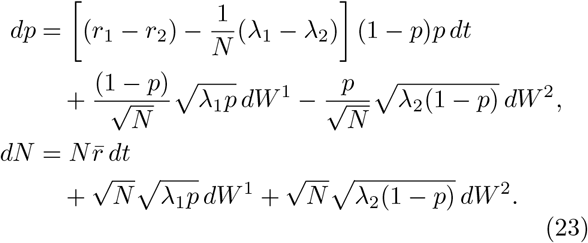

As we are only interested in the distribution properties of *p* and *N* we can though modify the equations above a bit. We can notice that the Fokker–Planck equation for the process does not explicitly depend on the form of the matrix

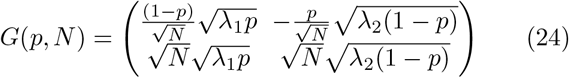

multiplying *dW* :s but instead, it is only function of the term

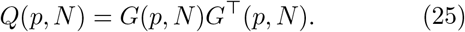

Therefore the distributional properties of *p* and *N* remain intact if we replace the matrix *G* with another matrix 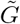 such that

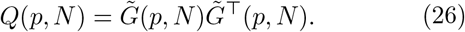

In our case we have

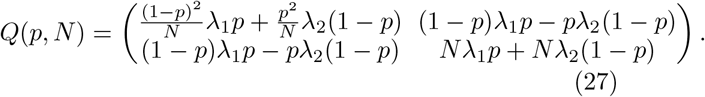

We can now select 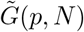 as the lower triangular Cholesky factorization of *Q*(*p, N*), that is,

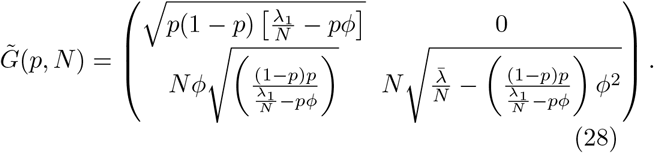

Thus if we introduce a 2D Wiener process 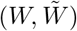 which has the same law as (*W*_1_, *W*_2_), we can form weakly equivalent SDEs for *p* and *N* as follows:

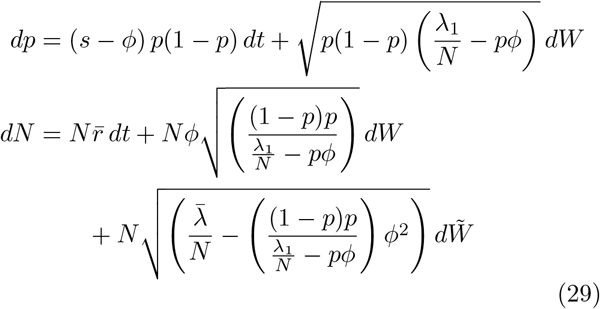

where *s* = *r*_1_ − *r*_2_ and 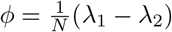.

In principle, we can also make use of a similar Cholesky factorization in general *d*-dimensional case, which allows for writing one of the probabilities (say *P*_*i*_) as function of a single Wiener process. However, all of them cannot be jointly written in such a form. The resulting equation for the single probability component can be written as

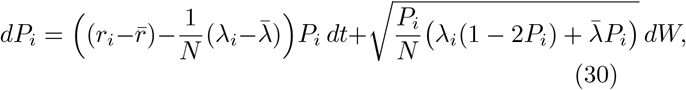

which indeed is a generalization of the first equation in (29).

### Patch model as a case study

In order to investigate mutant fixation, substitution and establishment we need to fix some ecological model that defines the effective birth and death rates. We note that the simplest and most widely used model, the logistic model, is unsuitable, because in all mechanistically derived instances of logistic growth the “carrying capacity” is not a constant, but rather a function of the intrinsic birth and death rates (for more discussion see e.g. [57]). Therefore assuming a constant carrying capacity term *K* while assuming the frequency dynamics of subtypes with different intrinsic rates is problematic. Hence we use a slightly modified version of the logistic model, which is more ecologically justified and also includes elements of frequency-dependent selection.

Consider the case where the growth is limited by the available space and let *K* denote the number of free patches, i.e. the maximum number of individuals the system can sustain (absolute carrying capacity which cannot be exceeded). Individuals die with their intrinsic death rate and produce a new offspring according to their intrinsic birth rate, which then randomly disperses to one of the patches. We can now consider two forms of density regulation depending on what happens if the new offspring lands on an occupied patch. There are only two cases: either the new offspring will die or replace and kill the the previous inhabitant. We can now consider a replacement probability *θ* ∈ [0, 1] (which we assume is determined by the ecological system and not the individual) that gives the probability that a birth on an occupied patch is successful.

With these rules set, the effective birth rate is the probability of landing on an empty patch 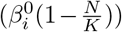 plus the probability of landing on an occupied patch times the replacement probability 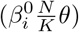. The effective death rate in turn is then the intrinsic death rate plus the deaths induced by the replacing offspring from both the own and competing strains. The total rate of producing offspring is given by 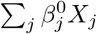, they land on any specific patch with probability 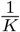 and replace with probability *θ*. Hence the product 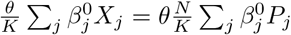 gives the death rate caused by replacement. This gives equations

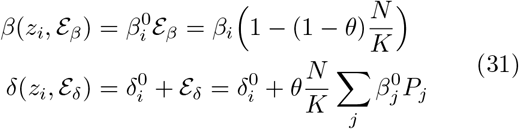

where the ecological feedback acts multiplicatively on the birth rate and additively on the death rate. These rates then determine the effective growth and turnover rates

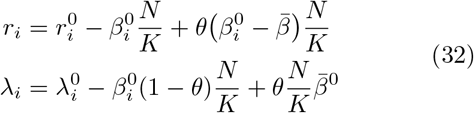

where 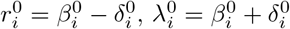, and 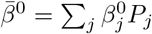.

#### Fixation in patch models

We can use the Kimura equation [22] to derive the fixation probabilities analytically, but taking care that we substitute the effective growth and turnover rates (32) when calculating the infinitesimal mean and variance.

First, let us calculate the selection coefficient and turnover flux

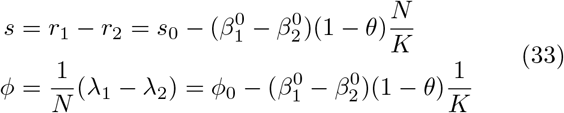

where we have defined 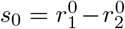 and 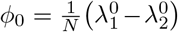. We note that the frequency-dependence in the death rates cancels out because the feedback ℰ_*δ*_ acts additively. Therefore, the selection coefficient and the turnover flux are purely density-dependent functions, which we can now substitute to (3). Thus, we find the infinitesimal mean

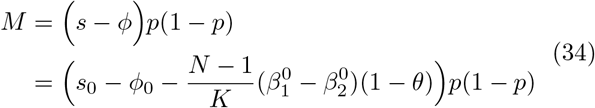

and variance

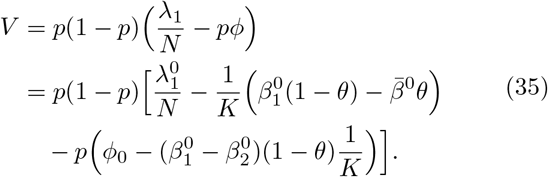

We note that these terms still contain additional time-dependencies via the *N* and 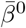 terms. To facilitate the analytical solution of the fixation probabilities, we resort to standard adaptive dynamics assumptions: first, it is easy to see that the resident’s population size equilibrates at

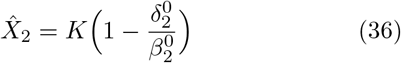

which allows us to set the initial condition 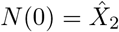 by assuming that the resident population equilibrates before the next mutant arrives. Next, we further approximate that 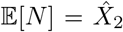 throughout the relevant stochastic dynamics during the mutant establishment, which in effect allows us to treat *N* as constant before the population size re-equilibrates to the mutant’s equilibrium 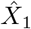 upon fixation. We can then take this value as the new initial condition for the subsequent substitution events. Finally, we neglect the minor frequency-dependency in the mean birth rate and approximate 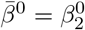.

Now, by defining the constants 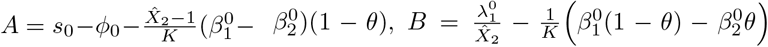 and 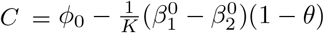, we get the integrator

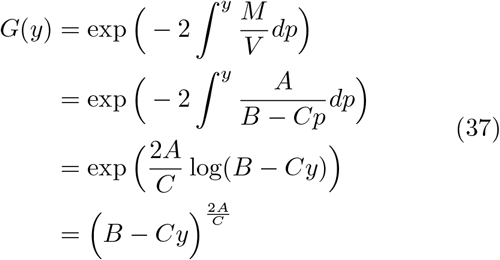

from which we can now obtain the fixation probability for a mutant starting from initial frequency *p* as

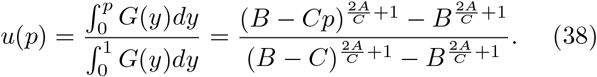

We can further study the pure birth-death process as a special case by removing the ecological feedback by taking the patch size *K* → ∞. This leads to the fixation probability given in (4).

### Establishment threshold

Let us consider the SDEs for *p* and *N* given in (29). Using Ito’s lemma (see, e.g., [36]) for function log *p* then gives

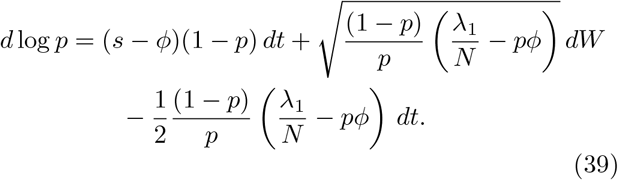

Now the value that forces the drift to be constant is given by

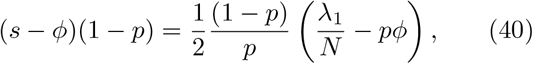

where we get that either *p*^*^ = 1 or then

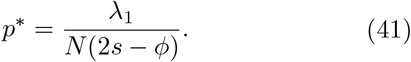

The same result would be obtained by using the log-transformation on Eqs. (23) instead. In the general *d*-dimensional case we can follow the same procedure by applying the log-transform to Eq. (30) and in principle recover a similar threshold equation in an implicit form.

### Stochastic simulations

All the stochastic simulations were done by simulating the discrete birth-death equation (17) forward in time using the patch model equations (31) while varying the intrinsic rates as well as the parameter *θ* which determines whether the ecological feedback is channeled via decreased birth (*θ* = 0) or increased death (*θ* = 1). Figure 2 had 10^6^ independent realisations per data point and Figure 3 10^3^ trajectories per initial mutant fraction. The evolutionary trajectories displayed in Fig. 5 were simulated starting from the initial trait *z*_0_ = (0.5, 0.3) and assuming an unbiased Gaussian mutation kernel 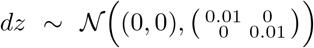 with per capita mutation rate *µ* = 0.005. Because this results in multiple coexisting traits, we sped up the simulations by first drawing the next generation size from a Poisson distribution 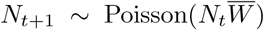 using mean multiplicative fitness 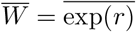 and then sampling the genotypes using weights *W*_*i*_ = exp(*r*_*i*_) (see e.g. [58]). The concave trade-off curve used for the feasible trait space was defined as *B*(*δ*_0_) = 0.6·log(*δ*_0_)+2.2−1.8·*δ*_0_ which gives the maximal intrinsic birth rate for given intrinsic death rate. The plotted mean evolutionary trajectory was calculated as the mean trait 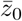 over 1000 simulated trajectories, each with length 10^4^ time steps.

## ACKNOWLEDGMENTS

We would like to thank Alexandre Minetto for comments on an earlier version of the work.

## AUTHOR CONTRIBUTIONS

(**T.K**.) Conceptualization, Formal analysis, Investigation, Methodology, Writing – original draft, Writing – review & editing. (**S.S**.) Formal analysis, Methodology, Writing – review & editing. (**V.M**.) Formal analysis, Investigation, Methodology, Supervision, Writing – review & editing.

## References

[1] D. L. Hartl and A. G. Clark, Principles of population genetics (4th ed.) (Oxford University Press, 2007).

[2] J. Berg, S. Willmann, and M. Lässig, Adaptive evolution of transcription factor binding sites, BMC evolutionary biology 4, 1 (2004).

[3] S. Aeschbacher, J. P. Selby, J. H. Willis, and G. Coop, Population-genomic inference of the strength and timing of selection against gene flow, Proceedings of the National Academy of Sciences 114, 7061 (2017).

[4] M. Gerstung, C. Jolly, I. Leshchiner, S. C. Dentro, S. Gonzalez, D. Rosebrock, T. J. Mitchell, Y. Rubanova, P. Anur, K. Yu, M. Tarabichi, A. Deshwar, J. Wintersinger, K. Kleinheinz, I. Vázquez-García, K. Haase, L. Jerman, S. Sengupta, G. Macintyre, S. Malikic, N. Donmez, D. G. Livitz, M. Cmero, J. Demeulemeester, S. Schumacher, Y. Fan, X. Yao, J. Lee, M. Schlesner, P. C. Boutros, D. D. Bowtell, H. Zhu, G. Getz, M. Imielinski, R. Beroukhim, S. C. Sahinalp, Y. Ji, M. Peifer, F. Markowetz, V. Mustonen, K. Yuan, W. Wang, Q. D. Morris, S. C. Dentro, I. Leshchiner, M. Gerstung, C. Jolly, K. Haase, M. Tarabichi, J. Wintersinger, A. G. Deshwar, K. Yu, S. Gonzalez, Y. Rubanova, G. Macintyre, D. J. Adams, P. Anur, R. Beroukhim, P. C. Boutros, D. D. Bowtell, P. J. Campbell, S. Cao, E. L. Christie, M. Cmero, Y. Cun, K. J. Dawson, J. Demeulemeester, N. Donmez, R. M. Drews, R. Eils, Y. Fan, M. Fittall, D. W. Garsed, G. Getz, G. Ha, M. Imielinski, L. Jerman, Y. Ji, K. Kleinheinz, J. Lee, H. Lee-Six, D. G. Livitz, S. Malikic, F. Markowetz, I. Martincorena, T. J. Mitchell, V. Mustonen, L. Oesper, M. Peifer, M. Peto, B. J. Raphael, D. Rosebrock, S. C. Sahinalp, A. Salcedo, M. Schlesner, S. Schumacher, S. Sengupta, R. Shi, S. J. Shin, O. Spiro, L. D. Stein, I. Vázquez-García, S. Vembu, D. A. Wheeler, T.-P. Yang, X. Yao, K. Yuan, H. Zhu, W. Wang, Q. D. Morris, P. T. Spellman, D. C. Wedge, P. Van Loo, P. T. Spellman, D. C. Wedge, P. Van Loo, P. E.. H. W. Group, and P. Consortium, The evolutionary history of 2,658 cancers, Nature 578, 122 (2020).

[5] P. A. Zur Wiesch, R. Kouyos, J. Engelstädter, R. R. Regoes, and S. Bonhoeffer, Population biological principles of drug-resistance evolution in infectious diseases, The Lancet infectious diseases 11, 236 (2011).

[6] H. Li and R. Durbin, Inference of human population history from individual whole-genome sequences, Nature 475, 493 (2011), 1011.1669v3.

[7] L. B. Alexandrov, S. Nik-Zainal, D. C. Wedge, S. A. Aparicio, S. Behjati, A. V. Biankin, G. R. Bignell, N. Bolli, A. Borg, A. L. Børresen-Dale, S. Boyault, B. Burkhardt, A. P. Butler, C. Caldas, H. R. Davies, C. Desmedt, R. Eils, J. E. Eyfjörd, J. A. Foekens, M. Greaves, F. Hosoda, B. Hutter, T. Ilicic, S. Imbeaud, M. Imielinsk, N. Jäger, D. T. Jones, D. Jonas, S. Knappskog, M. Koo, S. R. Lakhani, C. López-Otín, S. Martin, N. C. Munshi, H. Nakamura, P. A. Northcott, M. Pajic, E. Papaemmanuil, A. Paradiso, J. V. Pearson, X. S. Puente, K. Raine, M. Ramakrishna, A. L. Richardson, J. Richter, P. Rosenstiel, M. Schlesner, T. N. Schumacher, P. N. Span, J. W. Teague, Y. Totoki, A. N. Tutt, R. Valdés-Mas, M. M. Van Buuren, L. Van ‘T Veer, A. Vincent-Salomon, N. Waddell, L. R. Yates, J. Zucman-Rossi, P. Andrew Futreal, U. McDermott, P. Lichter, M. Meyerson, S. M. Grimmond, R. Siebert, E. Campo, T. Shibata, S. M. Pfister, P. J. Campbell, and M. R. Stratton, Signatures of mutational processes in human cancer, Nature 500, 415 (2013), NIHMS150003.

[8] M. Kimura, The neutral theory of molecular evolution (Cambridge University Press, 1983).

[9] M. Nei, The new mutation theory of phenotypic evolution, Proceedings of the National Academy of Sciences 104, 12235 (2007).

[10] A. M. Leroi, B. Lambert, J. Rosindell, X. Zhang, and G. D. Kokkoris, Neutral syndrome, Nature human behaviour 4, 780 (2020).

[11] M. A. McPeek, The ecological dynamics of natural selection: traits and the coevolution of community structure, The American Naturalist 189, E91 (2017).

[12] D. M. Fitzgerald and S. M. Rosenberg, What is mutation? a chapter in the series: How microbes “jeopardize” the modern synthesis, PLoS genetics 15, e1007995 (2019).

[13] M. Russo, A. Sogari, and A. Bardelli, Adaptive evolution: How bacteria and cancer cells survive stressful conditions and drug treatment, Cancer Discovery 11, 1886 (2021).

[14] D. W. Pfennig, Phenotypic plasticity & evolution: causes, consequences, controversies (Taylor & Francis, 2021).

[15] D. Nichol, M. Robertson-Tessi, A. R. Anderson, and P. Jeavons, Model genotype–phenotype mappings and the algorithmic structure of evolution, Journal of The Royal Society Interface 16, 20190332 (2019).

[16] G. W. Constable, T. Rogers, A. J. McKane, and C. E. Tarnita, Demographic noise can reverse the direction of deterministic selection, Proceedings of the National Academy of Sciences 113, E4745 (2016).

[17] D. A. Roff, Defining fitness in evolutionary models, Journal of genetics 87, 339 (2008).

[18] M. Doebeli, Y. Ispolatov, and B. Simon, Point of view: Towards a mechanistic foundation of evolutionary theory, Elife 6, e23804 (2017).

[19] A. G. Casanova, V. M. Pina, and J. C. Pardo, The wright–fisher model with efficiency, Theoretical Population Biology 132, 33 (2020).

[20] A. E. Noble, A. Hastings, and W. F. Fagan, Multivariate moran process with lotka-volterra phenomenology, Physical review letters 107, 228101 (2011).

[21] G. W. Constable and A. J. McKane, Mapping of the stochastic lotka-volterra model to models of population genetics and game theory, Physical Review E 96, 022416 (2017).

[22] M. Kimura, On the probability of fixation of mutant genes in a population, Genetics 47, 713 (1962).

[23] S. P. Otto and M. C. Whitlock, The probability of fixation in populations of changing size, Genetics 146, 723 (1997).

[24] Z. Patwa and L. M. Wahl, The fixation probability of beneficial mutations, Journal of The Royal Society Interface 5, 1279 (2008).

[25] P. Czuppon and A. Traulsen, Fixation probabilities in populations under demographic fluctuations, Journal of mathematical biology 77, 1233 (2018).

[26] H. Alexander and L. Wahl, Fixation probabilities depend on life history: fecundity, generation time and survival in a burst-death model, Evolution: International Journal of Organic Evolution 62, 1600 (2008).

[27] Y. Vindenes, A. M. Lee, S. Engen, and B.-E. Sæther, Fixation of slightly beneficial mutations: effects of life history, Evolution: International Journal of Organic Evolution 64, 1063 (2010).

[28] X.-Y. Li, S. Kurokawa, S. Giaimo, and A. Traulsen, How life history can sway the fixation probability of mutants, Genetics 203, 1297 (2016).

[29] D. Abu Awad and C. Coron, Effects of demographic stochasticity and life-history strategies on times and probabilities to fixation, Heredity 121, 374 (2018).

[30] J. E. Hubbarde, G. Wild, and L. M. Wahl, Fixation probabilities when generation times are variable: The burst– death model, Genetics 176, 1703 (2007).

[31] T. L. Parsons, C. Quince, and J. B. Plotkin, Some consequences of demographic stochasticity in population genetics, Genetics 185, 1345 (2010).

[32] P. Ashcroft, A. Traulsen, and T. Galla, When the mean is not enough: Calculating fixation time distributions in birth-death processes, Physical Review E 92, 042154 (2015).

[33] P. Czuppon and A. Traulsen, Understanding evolutionary and ecological dynamics using a continuum limit, Ecology and Evolution 11, 5857 (2021).

[34] V. Mustonen and M. Lässig, From fitness landscapes to seascapes: non-equilibrium dynamics of selection and adaptation, Trends in Genetics 25, 111 (2009).

[35] T. L. Vincent and J. S. Brown, Evolutionary game theory, natural selection, and Darwinian dynamics (Cambridge University Press, 2005).

[36] S. Särkkä and A. Solin, Applied stochastic differential equations, vol. 10 (Cambridge University Press, 2019).

[37] M. A. Nowak, Evolutionary dynamics: exploring the equations of life (Harvard university press, 2006).

[38] V. Mustonen and M. Lassig, Adaptations to fluctuating selection in Drosophila, Proc Natl Acad Sci U S A. 104, 2277 (2007).

[39] I. M. Rouzine, A. Rodrigo, and J. Coffin, Transition between stochastic evolution and deterministic evolution in the presence of selection: general theory and application to virology, Microbiology and molecular biology reviews 65, 151 (2001).

[40] M. M. Desai and D. S. Fisher, Beneficial mutation– selection balance and the effect of linkage on positive selection, Genetics 176, 1759 (2007).

[41] O. Peters, Optimal leverage from non-ergodicity, Quantitative Finance 11, 1593 (2011).

[42] O. Peters and W. Klein, Ergodicity breaking in geometric brownian motion, Physical review letters 110, 100603 (2013).

[43] J. Lehtonen, The price equation, gradient dynamics, and continuous trait game theory, The American naturalist 191, 146 (2018).

[44] J. C. Baez, The fundamental theorem of natural selection, Entropy 23, 1436 (2021).

[45] M. Gyllenberg and K. Parvinen, Necessary and sufficient conditions for evolutionary suicide, Bulletin of mathematical biology 63, 981 (2001).

[46] R. Sear, D. W. Lawson, H. Kaplan, and M. K. Shenk, Understanding variation in human fertility: what can we learn from evolutionary demography?, Philosophical Transactions of the Royal Society B: Biological Sciences 371, 20150144 (2016).

[47] J.-B. André and F. Rousset, Does extrinsic mortality accelerate the pace of life? a bare-bones approach, Evolution and Human Behavior 41, 486 (2020).

[48] C. de de Vries, M. Galipaud, and H. Kokko, Extrinsic mortality and senescence: a guide for the perplexed, bioRxiv (2022).

[49] J. Rybicki, E. Kisdi, and J. V. Anttila, Model of bacterial toxin-dependent pathogenesis explains infective dose, Proceedings of the National Academy of Sciences 115, 10690 (2018).

[50] D. J. Hruschka and O. Burger, How does variance in fertility change over the demographic transition?, Philosophical Transactions of the Royal Society B: Biological Sciences 371, 20150155 (2016).

[51] C. A. Aktipis, A. M. Boddy, R. A. Gatenby, J. S. Brown, and C. C. Maley, Life history trade-offs in cancer evolution, Nature Reviews Cancer 13, 883 (2013).

[52] J. A. Gallaher, J. S. Brown, and A. R. Anderson, The impact of proliferation-migration tradeoffs on phenotypic evolution in cancer, Scientific reports 9, 1 (2019).

[53] J. V. Anttila, M. Shubin, J. Cairns, F. Borse, Q. Guo, T. Mononen, I. Vázquez-García, O. Pulkkinen, and V. Mustonen, Contrasting the impact of cytotoxic and cytostatic drug therapies on tumour progression, PLoS computational biology 15, e1007493 (2019).

[54] T. Kuosmanen, J. Cairns, R. Noble, N. Beerenwinkel, T. Mononen, and V. Mustonen, Drug-induced resistance evolution necessitates less aggressive treatment, PLoS computational biology 17, e1009418 (2021).

[55] R. Sender and R. Milo, The distribution of cellular turnover in the human body, Nature Medicine 27, 45 (2021).

[56] A. Mazzolini and J. Grilli, Universality of evolutionary trajectories under arbitrary competition dynamics, bioRxiv 10.1101/2021.06.17.448795 (2022), https://www.biorxiv.org/content/early/2022/05/06/2021.06.17.448795.full.pdf.

[57] J. Mallet, The struggle for existence. how the notion of carrying capacity, k, obscures the links between demography, darwinian evolution and speciation, Evolutionary Ecology Research (2012).

[58] Y. Anciaux, L.-M. Chevin, O. Ronce, and G. Martin, Evolutionary rescue over a fitness landscape, Genetics 209, 265 (2018).

